# Dissecting MPXV neurotropism and host antiviral signaling using human stem cell-based models

**DOI:** 10.1101/2023.08.25.554849

**Authors:** Lisa Bauer, Stefania Giussani, Nicola Palazzi, Erik Bot, Elisa Colombo, Farnaz Zare, Francesca Pinci, Lonneke Leijten, Kristina Lanko, Feline F.W. Benavides, Hilde Smeenk, Carmen W.E. Embregts, Jochem K.H. Spoor, Clemens Dirven, Zhenyu Gao, Anne Bolleboom, Babs E. Verstrepen, Leonard Schuele, Femke M.S. de Vrij, Steven A. Kushner, Bas B. Oude Munnink, Jose Davila-Velderrain, Debby van Riel, Oliver Harschnitz

## Abstract

Mpox is a zoonotic illness of international concern that can lead to severe disease including neurological sequelae. However, the neurotropism of monkeypox virus (MPXV) and the mechanisms regulating cell-intrinsic antiviral immunity within the central nervous system (CNS) remain poorly understood. Here, we investigated the neurotropism of MPXV using astrocytes, cortical neurons, and microglia derived from human pluripotent stem cells (hPSCs) and *ex vivo* human brain tissue to demonstrate that MPXV infects and replicates more efficiently in astrocytes and microglia compared to cortical neurons. Upon MPXV exposure, glial cells, in contrast to cortical neurons, inhibit type I IFN antiviral programs potentially conferring differential susceptibility to MPXV. Furthermore, we demonstrate that treatment using either IFN-beta or tecovirimat inhibits MPXV infection. Together, our results suggest that MPXV has a broad tropism within the CNS and that differential type I IFN signaling underpins cell type-specific susceptibility to MPXV infection.

## Introduction

Mpox, formerly known as monkeypox(Ulaeto et al., 2023), was declared a global public health emergency of international concern by the World Health Organization twice in quick succession, the first time in July 2022 after a multi-country outbreak of clade IIb and the second time in August 2024 as cases with clade Ia and clade Ib have continued to increase and spread outside areas where it has traditionally been endemic(Thornhill et al., 2022a; de Vries et al., 2023; Nzoyikorera et al., 2024). The threat of monkeypox virus (MPXV) and other orthopoxviruses is high due to decreasing levels of population immunity after cessation of smallpox vaccination, as well as the overall lack of knowledge on poxvirus cross-species transmission and the pathogenesis on a host and cellular level(McFadden, 2005; Rimoin et al., 2010; Lu et al., 2023). Mpox patients typically present with a characteristic rash with further systemic manifestations including fever, fatigue, and myalgia(Fink et al., 2022; Thornhill et al., 2022b). However, complications include sepsis, pneumonia, conjunctivitis, and neurological sequelae and severe cases can be fatal. In the majority of severe cases, patients present with neuropsychiatric complications including seizures, confusion, and encephalitis, indicating that MPXV is able to invade the central nervous system (CNS)(Fink et al., 2022; Badenoch et al., 2022; Cole et al., 2022; Billioux et al., 2022; Pastula and Tyler, 2022; Sharma et al., 2023). Recent studies using primary human brain tissue and cells derived from human pluripotent stem cells (hPSCs) have suggested a tropism of MPXV for human CNS cells(Chailangkarn et al., 2022; Mahabadi et al., 2024). However, the cell type-specific neurotropism of MPXV within the brain has remained controversial(Schultz-Pernice et al., 2023) and the molecular mechanisms underlying cell type-specific antiviral immunity within the CNS are poorly understood, hampering the discovery of potential therapeutic targets for the neuropsychiatric complications of this emerging zoonotic disease.

As brain tissue from infected patients is scarce and can only provide insights into the end-phase of the disease, *in vitro* models are necessary to study MPXV neurotropism and the underlying antiviral mechanisms in human brain cells. Human PSC-based models have emerged as powerful tools for studying host-virus interactions of the CNS *in vitro*(Harschnitz and Studer, 2021). Recent studies applying hPSC-derived models have significantly enhanced our understanding of the neurotropism and neurovirulence of newly emerging pathogens, such as SARS-CoV-2(Pellegrini et al., 2020; Jacob et al., 2020; Yang et al., 2020, 2024; Chan et al., 2024). Hence, we used a hPSC-based platform to systematically dissect the neurotropism of clade IIb MPXV in human, disease-relevant CNS cells. Our results suggest a broad tropism of MPXV within the CNS, with a preferential tropism for glial cells over cortical neurons supporting the notion that not all brain cells are equally susceptible to MPXV infection. We identified distinct molecular changes in neurons and glia, where the latter downregulate their type I IFN antiviral signaling upon MPXV exposure. Lastly, we show that MPXV CNS infection could be ameliorated using antiviral treatment with either IFN-beta or tecovirimat.

## Results

### Differential susceptibility of human CNS cells to MPXV infection

Human PSCs can be differentiated into all major brain cell lineages, and recapitulate key aspects of CNS function and pathophysiology(Grant Rowe and Daley, 2019; Harschnitz and Studer, 2021). To study the neurotropism of MPXV, we performed directed differentiation of hPSCs (WA-09) into pure populations of astrocytes(Lendemeijer et al., 2024) (**Fig. S1a and 1b**), cortical neurons(Ciceri et al., 2022) (**Fig. S1c-h**), and microglia(Guttikonda et al., 2021) (**Fig. S1i-l**) using previously reported protocols. To determine the relative susceptibility, permissiveness, and response to infection, we inoculated these human CNS cells with cell-culture propagated clade IIb MPXV at a multiplicity of infection (MOI) of 1 (**Fig. 1a-k**). Immunofluorescence staining for MPXV revealed that astrocytes and microglia were considerably more susceptible to MPXV infection than cortical neurons, which increased over time, suggesting that not all CNS populations are equally susceptible to MPXV infection (**Fig. 1a, 1b, 1d, 1e, 1g, and 1h**). Furthermore, we observed pronounced cell death in hPSC-derived astrocytes exposed to MPXV from 48 hours onwards (**Fig. 1a**). This recapitulates what has previously been reported using human primary astrocytes, which undergo pyroptosis-mediated cell death following MPXV infection(Mahabadi et al., 2024). No MPXV antigen was detected in mock-infected cell cultures (**Fig. 1a, 1d, and 1g**).

**Fig. 1:**
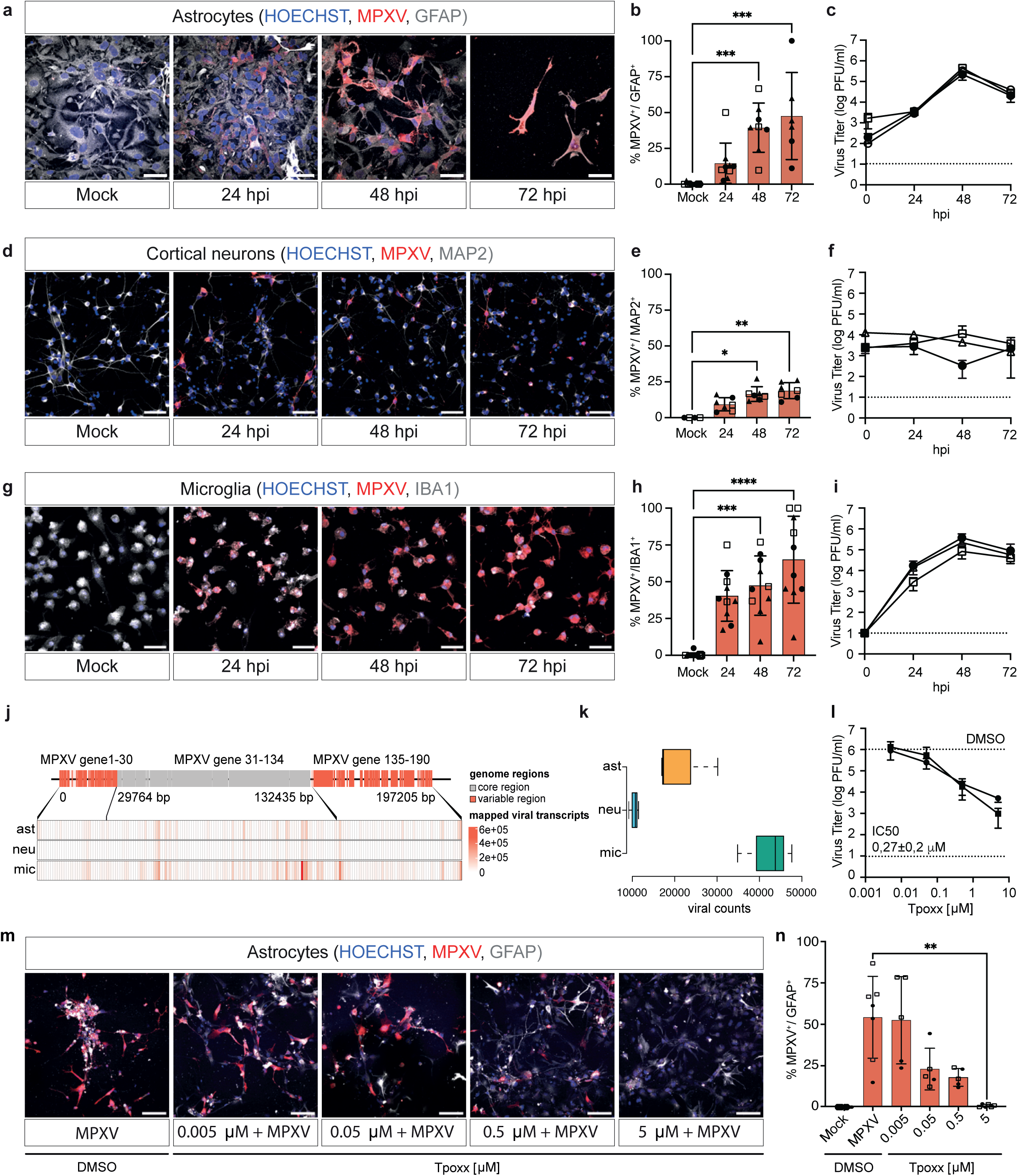
Astrocytes, cortical neurons, and microglia are differentially susceptible and permissive to MPXV infection. **a-b,** Immunofluorescence staining for astrocyte protein GFAP (grey) and MPXV antigen (red), with quantification (**b**) of MPXV+/GFAP+ cells. Images representative of three independent experiments. **c,** Plaque forming assays of supernatant collected from MPXV-infected astrocytes. **d-e,** Immunofluorescence staining for neuronal protein MAP2 (grey) and MPXV antigen (red) with quantification (**e**) of MPXV+/MAP2+ cells. Images representative of three independent experiments. **f,** Plaque forming assays of supernatant collected from MPXV-infected cortical neurons. **g-h,** Immunofluorescence staining for microglial protein IBA1 (grey) and MPXV antigen (red) with quantification (**h**) of MPXV+/IBA1+ cells. Images representative of three independent experiments. **i,** Plaque forming assays of supernatant collected from MPXV-infected microglia. **j,** Alignment and distribution of viral reads to the viral genome per cell-type. **k,** Normalized viral reads following RNA-seq 24 hours after MPXV infection of astrocytes, cortical neurons, and microglia. **l,** In a multi-cycle replication assay, astrocytes were infected with MPXV (MOI = 1) and treated with a serial dilution of Tpoxx for 48 hrs. Plaque forming assay was used to determine MPXV titers upon Tpoxx treatment. IC50 is indicated. **m**, Immunofluorescence staining of astrocytes (GFAP+, grey) and MPXV antigen (red) 48 hours post infection. **n,** Quantification of MPXV^+^/GFAP^+^ infected astrocytes upon Tpoxx treatment. **a-i,** *N* = 3 independent biological replicates, indicated by different shapes. *n* = 7-10 technical replicates. **l-n,** *N* = 2 independent biological replicates, *n* = 5-7 technical replicates. **a-k**, Experiments were performed using WA-09-derived astrocytes, cortical neurons and microglia. **l-n**, Experiments were performed using WTC11-derived astrocytes. Statistical analysis was performed using Kruskal-Wallis one-way ANOVA with Dunn’s correction for multiple comparisons. *p < 0.05, **p < 0.01, ***p < 0.001, ****p < 0.0001. All data represent mean ± SD from pooled experiments. All experiments were performed in two to three biological replicates.

### MPXV productively infects glial cells

To determine the replication efficiency, we collected supernatant daily following MPXV inoculation and performed plaque forming assays on Vero E6 cells (**Fig. S2a-S2c**). This analysis confirmed productive infection in human astrocytes and microglia, based on an increase in viral titers over time (**Fig. 1c and 1i**). No significant increase in viral titer was observed in cortical neurons (**Fig. 1f**). RNA sequencing (RNA-seq) confirmed differences in viral replication efficiency among CNS cells, with the lowest number of viral transcripts detected in cortical neurons compared to astrocytes and microglia at 24 hpi (**Fig. 1j and 1k**).

To further validate the susceptibility and permissiveness of human CNS cells to MPXV infection, we derived astrocytes, cortical neurons, and microglia from an additional hPSC line (WTC-11(Kreitzer et al., 2013)). Immunostaining (**Fig. S2d, S2e, S2j, and S2k)** and plaque forming assays (**Fig. S2f and S2l)** confirmed robust MPXV infection in WTC-11 hPSC-derived astrocytes and microglia, in contrast to cortical neurons (**Fig. S2g-S2i)**. Together, these results demonstrate that hPSC-derived astrocytes and microglia, and to a lesser extent cortical neurons, are susceptible and permissive to MPXV infection.

### Tecovirimat treatment inhibits MPXV replication

Drug sensitivity of smallpox antivirals, such as tecovirimat (Tpoxx), has been tested for the circulating MPXV clades in cell lines and primary cultures, such as human foreskin fibroblasts, human foreskin keratinocytes, and kidney organoids(Frenois-Veyrat et al., 2022; Warner et al., 2022; Bojkova et al., 2022; Li et al., 2023a). The efficacy of antivirals is highly dependent on the model and cell-type used(Bojkova et al., 2022), and the use of Tpoxx has not been tested in CNS cells. To test the efficacy of Tpoxx to treat mpox-induced neuropathology, we inoculated hPSC-derived astrocytes (the most severely affected CNS cell-type tested, see **Fig. 1**) with MPXV (MOI = 1), treated cells with serial dilutions of Tpoxx (0.005-5 µM), and tested for antiviral efficacy (**Fig. 1l-n**). Tpoxx treatment led to a significant dose-dependent reduction of viral titers at 48 hpi (**Fig. 1l**), in line with the mechanism-of-action of tecovirimat, which specifically prevents the production of infectious progeny by inhibiting the VP37 envelope wrapping protein(Yang et al., 2005; Grosenbach et al., 2011). Moreover, Tpoxx treatment also led to a dose-dependent reduction in MPXV-infected astrocytes in a multi-cycle infection assay and prevented MPXV-induced cell death up to 48 hpi (**Fig. 1m and 1n**). Together, these data suggest that Tpoxx treatment inhibits viral spread and the associated MPXV-induced cell death in human astroglial cells and could therefore be a potential therapeutic candidate to treat mpox encephalitis.

### Primary human brain tissue is susceptible and permissive to MPXV infection

To further confirm that human CNS cells are susceptible to MPXV infection, we inoculated primary *ex vivo* human post-surgical neocortical tissue with MPXV (6*10^6^ PFU) (**Fig. 2a**). We observed an increase in viral titers at 24 hpi, confirming primary human brain cells are both susceptible and permissive to MPXV infection (**Fig. 2b**). MPXV infection in human astrocytes and microglia was observed by co-expression of cell-types specific markers (GFAP and IBA1 immunofluorescence staining, respectively) and RNAscope for MPXV-specific RNA(Li et al., 2023b) at 24 hpi (**Fig. 2c-2e**). Interestingly, only in rare cases did we observe any HuC/HuD+ neurons positive for MPXV (**Fig. 2d**), in line with our data obtained from hPSC-derived CNS cells (**Fig. 1**). No specific signal was detected by RNAscope for MPXV RNA in mock-infected tissue (**Fig. 2c-2e**). These data provide further evidence for the cellular tropism of MPXV within the human brain.

**Fig. 2:**
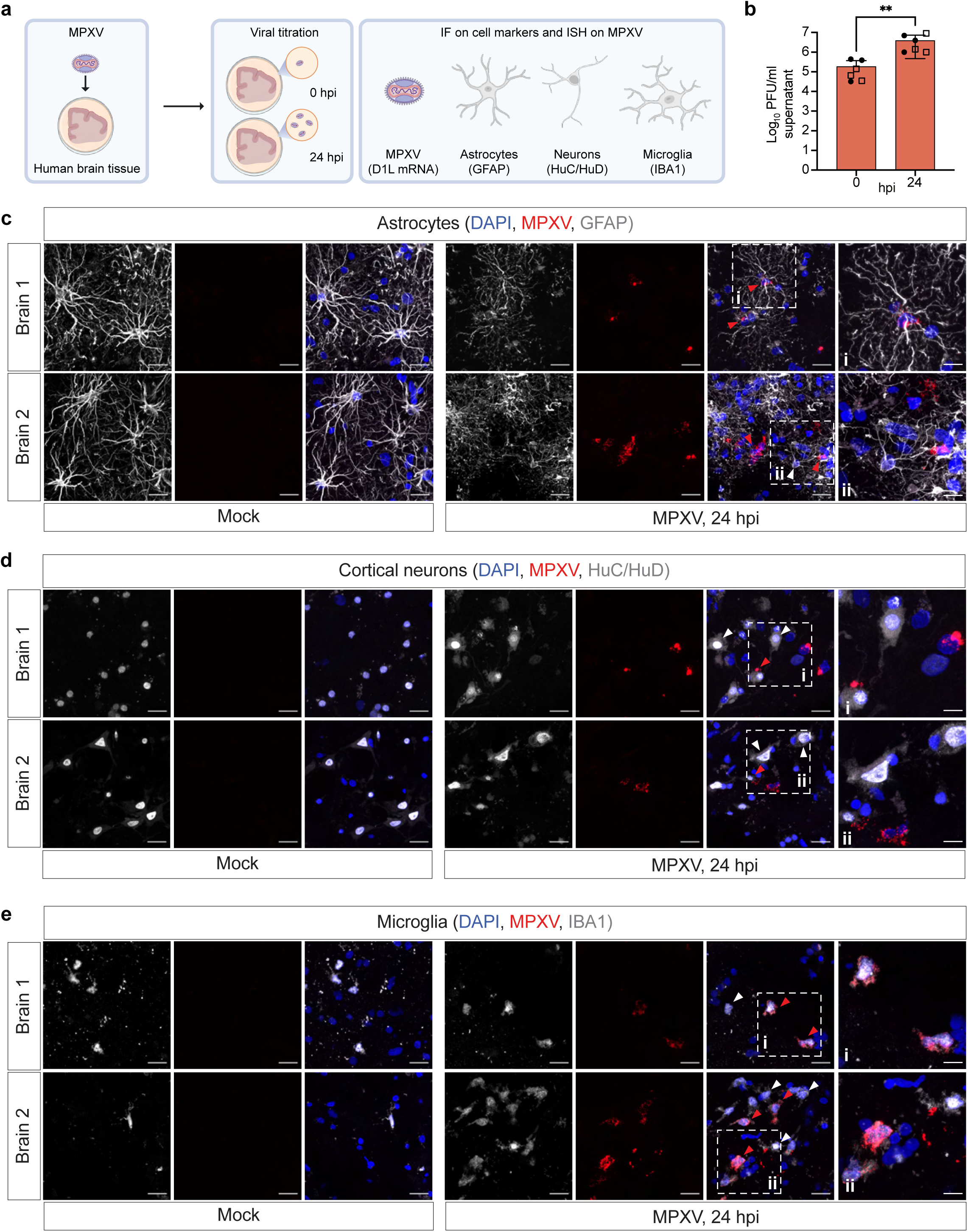
MPXV infects and replicates in *ex vivo* brain tissue. **a,** Schematics of human *ex vivo* brain tissue infection with MPXV (6x10^6 PFU), after which supernatant was collected for viral titration and a MPXV RNAscope was used to visualize the virus. Cells specific markers were detected by immunofluorescence staining. **b,** Plaque forming to determine MPXV titer in human *ex vivo* brain tissue 24 hpi. **c-e,** Representative images of mock- and MPXV-infected human brain slices stained for astrocyte marker GFAP, microglia marker IBA1, and neuronal marker HuC/HuD with fluorescent RNAscope to detect MPXV infection (red). Infected cells are indicated by red arrowheads, non-infected cells are indicated by white arrowheads. Scale bar = 20 µm. Scale bar of panels i and ii = 10 µm. **b**, *N*=2 independent biological replicates, *n* = 18 technical replicates. **c-e**, Stainings were performed on two independent brain tissues. Statistical analysis was performed using Mann Whitney t test, **p < 0.01.

### Transcriptomic changes in MPXV infected CNS cells

To determine the cell type-specific responses of human brain cells to MPXV infection, we performed RNA-seq of MPXV (MOI = 1, 24 hpi) and mock-infected hPSC-derived astrocytes, cortical neurons, and microglia. Human PSC-derivatives preferentially expressed known marker genes of astrocytes, cortical neurons, or microglia (**Fig. S1m**) consistent with the transcriptome profiles of their *in vivo* counterparts(Zhang et al., 2016) (**Fig. S1n**), further confirming their cellular identity. Consistent with the observed cell type-specific tropism (**Fig. 1**), transcriptomic responses to MPXV infection was most evident in astrocytes and microglia compared to cortical neurons (**Fig. S3a**), with astrocytes having a more variable within-sample response (**Fig. 3a**). To dissect the molecular basis of these changes, we performed differential expression analysis comparing mock versus infected samples for each cell type (**Fig. 3b and 3c**). Infected microglia showed the largest number of differentially regulated genes with a total of 2022 up-regulated and 1048 down-regulated genes (log2 fold change ≥ |1.0|, adjusted P < 0.01), of which 2584 were exclusive to this cell type (**Fig. 3b and 3c**). This was considerably more than in either cortical neurons (630 up-regulated and 275 down-regulated genes) or astrocytes (431 up-regulated and 92 down-regulated genes, **Fig. 3b**). More in-depth analyses revealed distinct differential gene expression patterns among the three cell types, with a subset of genes unique to each cell type (**Fig. 3d**), some differentially expressed genes shared by all cell types (**Fig. 3e**), and some shared by at least two cell types or some with opposite direction in any cell type pair (**Fig. S3b and S3c**). Other genes showed downregulation only in glial cell types (**Fig. 3f**). Of note, we also identified a subset of genes that was up-regulated in cortical neurons and down-regulated in glial cells, such as *SIGLEC1*, *SLC40A1* and *CYTH4*, suggesting a potential role for these genes in neuronal antiviral signaling (**Fig. 3f**). We then analyzed the differential pathway enrichment in all cell types (**Fig. 3g and S3d**). Genes that were found to be down-regulated in glial cell types were involved in immune-related pathways such as “Interferon alpha/beta signaling”, “COVID-19 disease”, and “Hepatitis C” (**Fig. 3g**). Interestingly, we found that the majority of downregulated pathways in astrocytes and, to a lesser extent, in microglia could be grouped under two macro-categories, namely “response to pathogens” and “immune response”. None of the top downregulated pathways in cortical neurons were related to antiviral immunity (**Fig. 3g**).

**Fig. 3:**
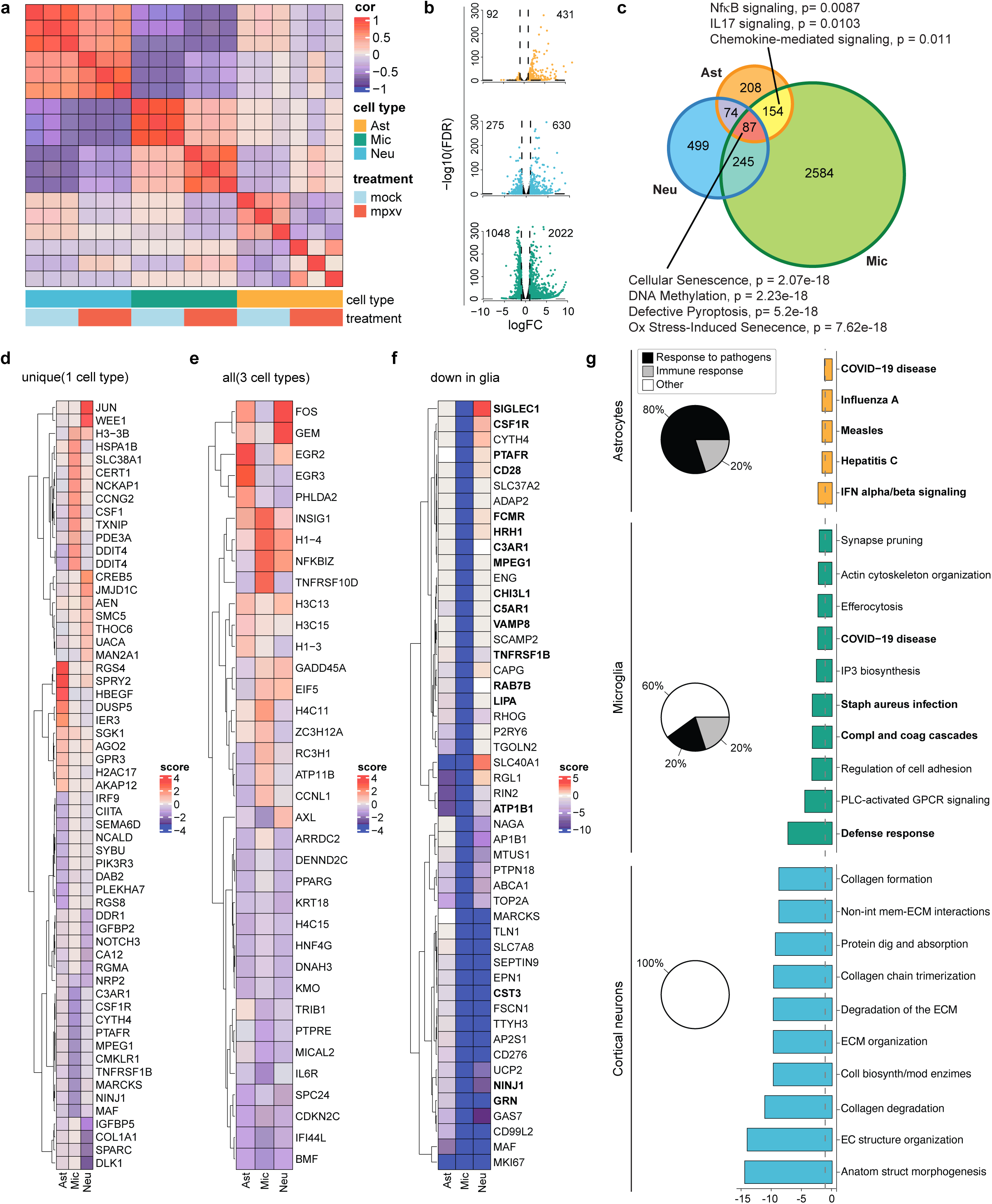
MPXV-infected glial cells downregulate immune-associated pathways. **a,** Pairwise correlations of transcriptional signatures (samplewise relative expression profiles). **b**, Volcano plot depicting differential expression analysis of mock- vs MPXV-infected hPSC-derived astrocytes, microglia, and cortical neurons. Number of differentially expressed genes (p-adjusted value < 0.01) with a log2 (MPXV/mock fold Change) ≥ |1.0| listed in top corner and color coded per cell type. **c**, Venn diagram depicting differentially expressed gene overlap among cell types. The area of the circles is in proportion to the size of the gene sets. Pathways with differential activity of specific gene sets from the Venn diagram are highlighted. **d-e**, Relative expression profiles for genes with a unique expression pattern, or one that is differentially expressed among all cell types. **f**, Expression pattern of genes that are downregulated in glial cells. Genes found at the intersection of those that are downregulated in the astrocytes and microglia and that belong to the pathways bolded in Figure 3G, are highlighted. **g,** Downregulated pathways in astrocytes, microglia and cortical neurons after MPXV infection with pie charts showing the proportion of pathways that are linked either to a response to pathogens (black) or to an immune response (grey). **a-g**, experiments were performed using WA-09-derived astrocytes, cortical neurons and microglia.

### MPXV infection inhibits innate antiviral immune response in glial cells

Host cells promote an antiviral state by producing and responding to type I interferon (IFN) through signal transducer and activator of transcription 1 (STAT1)-dependent signaling cascades(Ivashkiv and Donlin, 2014; McNab et al., 2015). Poxviruses, including vaccinia virus (VACV), variola virus (VARV) and MPXV, have the potential to evade the innate immune type I IFN system(Haga and Bowie, 2005; Fernández de Marco et al., 2010; Arndt et al., 2015; Talbot-Cooper et al., 2022). To characterize the specific effects of MPXV infection on antiviral-related processes within the CNS, we performed a targeted pathway analysis focusing on Gene Ontology biological processes involved in innate antiviral immunity. We aggregated gene expression values into pathway activity scores and estimated the effect of viral infection on these aggregate scores to perform differential pathway activity analysis (see Material and Methods). In homeostatic conditions, most pathways exhibited cell type-specific activity, yet a small subset was active across all three cell types (**Fig. 4a and 4b**). In line with cell type-specific phenotypes (**Fig. 1**), MPXV-infected astrocytes and microglia displayed a significant down-regulation for innate immune pathway activities (**Fig. 4a and 4b**). Notably, glial cells, in contrast to cortical neurons showed a decrease in type I IFN signaling (**Fig. 4a and 4b**). We validated these findings in astrocytes, cortical neurons, and microglia derived from a second hPSC line (WTC-11; **Fig. S4a and S4b**). These results suggest brain cell type-specific transcriptional responses to MPXV infection involving innate immunity may play a role in the increased susceptibility and permissiveness of astrocytes and microglia to MPXV infection and replication.

**Fig. 4:**
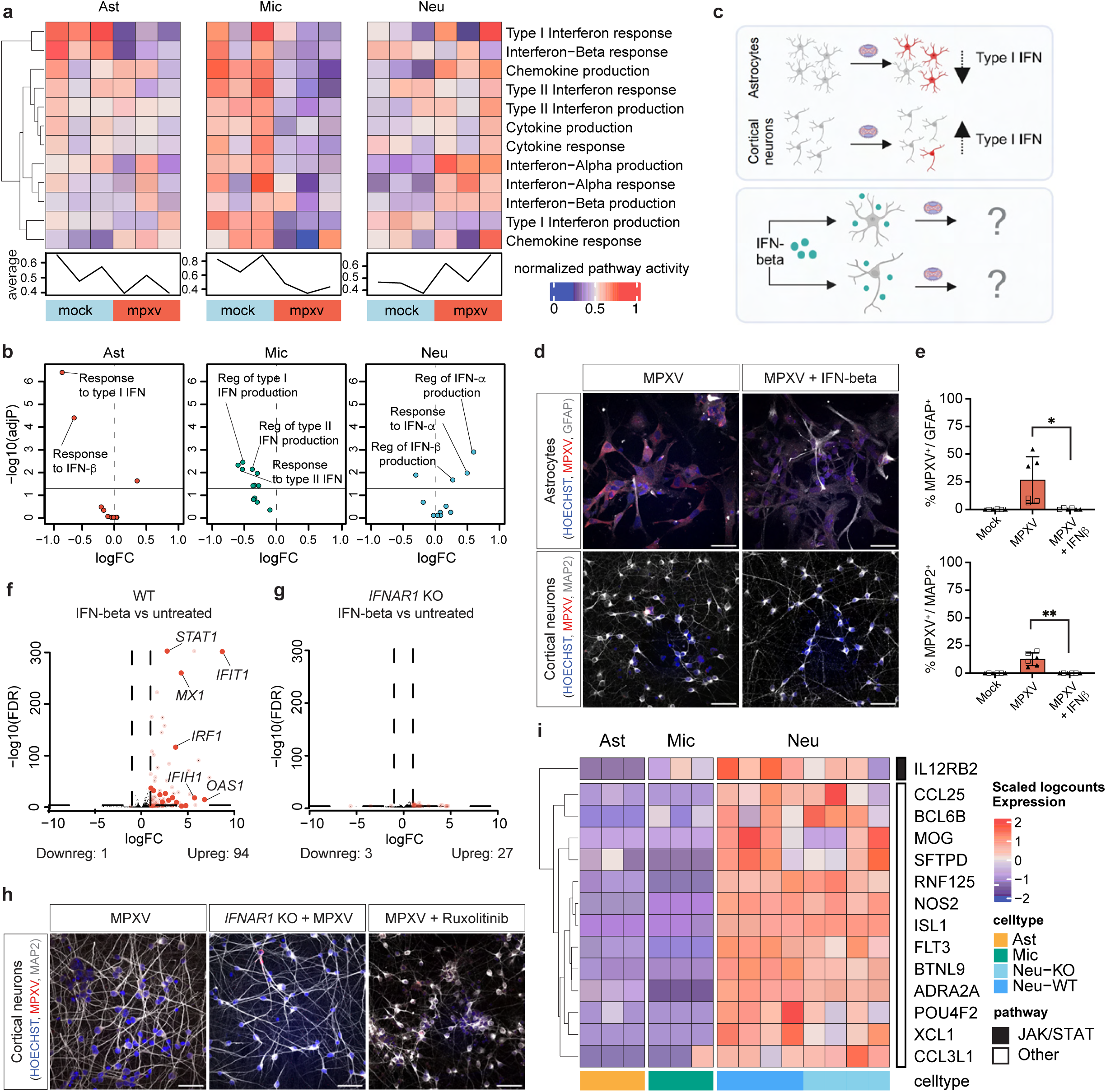
Type I Interferon signaling protects astrocytes and cortical neurons from MPXV infection. **a**, Pathway activity scores for interferon-associated GO processes. Scores were computed per sample within each cell type. Bottom line plot depicts average per sample activity of all antiviral processes (rows). **b**, Volcano plot depicting differential pathway activity analysis of interferon-associated GO processes in mock- vs MPXV-infected hPSC-derived astrocytes, microglia, and cortical neurons (adjusted p < 0.05, linear model, mock- versus MPXV-infected). **c**, Schematics representing the differential regulation of interferon signaling in glial cells and neurons upon infection. Experimental design to assess the protective effect of interferon-beta on astrocytes and cortical neurons. **d-e**, Immunofluorescence staining for astrocyte marker GFAP (upper panel) and neuronal protein MAP2 (lower panel) and MPXV antigen with quantification (**e**) of GFAP+/MPXV+ and MPXV+/MAP2+ cells respectively. **f-g**, Volcano plot depicting differential expression analysis of IFN-beta-treated vs untreated days *in vitro* (DIV)40 cortical neurons in the wt or *IFNAR1* KO background. The bold red dots indicate Interferon Stimulated Genes (ISGs) and some have been annotated in the plot. Number of differentially expressed genes (p-adjusted value < 0.01) with a log2 (IFN-beta treated/untreated fold Change) ≥ |1.0| listed below each plot. **h**, Immunofluorescence staining for MPXV-infected cortical neurons at 24 hpi. Images show, from left to right, wild type cortical neurons, *IFNAR1* KO cortical neurons and wild type cortical neurons pre-treated with Ruxolitinib, a JAK-STAT pathway inhibitor. **i**, Heatmap showing the baseline levels of genes belonging to antiviral pathways that are over-expressed between neurons (both wild type or *IFNAR1* KO) vs astrocytes and microglia (adjusted p-value < 0.05, logFC > 0.5). Black and white boxes represent genes that are respectively part or not of the JAK-STAT pathway. **e**, Experiments were performed using WTC11-derived astrocytes and WA-09-derived cortical neurons. **a, b, h, i,** Experiments were performed using WA-09-derived cortical neurons. **e**, *N* = 2 independent biological replicates for each cell type indicated by different shape of the data point, *n* = 6 technical replicates for cell type and condition. Statistical analysis of image quantifications was performed using one-way ANOVA with Kruskal Wallis test. *p < 0.05, **p < 0.01, ***p < 0.001, ****p < 0.0001. All data represent mean ± SD. All experiments were performed in two to three biological replicates

### Interferon treatment prevents MPXV infection

No specific treatment is currently approved for mpox encephalitis. Tecovirimat treatment rescues MPXV infection *in vitro* in hPSC-derived astrocytes (**Fig. 1l and 1m**) but may not be sufficient in a clinical setting. Indeed, recent studies in endemic regions of mpox showed no benefit on mortality following Tpoxx treatment in severe mpox cases (Clinical Trial n° NCT05559099) and the emergence of tecovirimat-resistant strains(Smith et al., 2023; Karan et al., 2024; Bapolisi et al., 2024; Akiyama et al., 2024). IFNs have been used as potential treatment for viral encephalitis, either as monotherapy or as combination therapy with other antivirals, however this is yet to be explored for poxvirus-induced encephalitis(Kalil et al., 2005; Wintergerst and Belohradsky, 1992). Previous reports demonstrate that MPXV production and spread in primary cell lines could be blocked by IFN-beta treatment(Johnston et al., 2012). Building on our data that suggest a key role for type I IFN signaling in cell-intrinsic anti-MPXV immunity in human brain cells (**Fig. 4a and 4b**), we treated hPSC-derived astrocytes, the most affected glial cell-type, and neurons with IFN-beta before inoculating cells with MPXV (MOI = 1) and tested for antiviral efficacy (**Fig. 4c**). Compared to DMSO-treated cells, IFN-beta treatment led to a reduction of MPXV-infected cells in both cell types (**Fig. 4d and 4e**), demonstrating an antiviral effect of type I IFNs against MPXV infection in human brain cells. These data suggest that type I IFN treatment is sufficient to boost anti-MPXV immunity in human CNS cells.

### Cortical neurons lacking IFNAR1 do not show increased susceptibility to MPXV infection

Next, we assessed whether inducible type I IFN signaling is essential for neuronal anti-MPXV immunity. Accordingly, to understand if homeostatic antiviral signaling or the response to type I IFNs was responsible for cortical neuron intrinsic immunity to MPXV infection, we generated *IFNAR1* knockout (KO) hPSCs and differentiated these into cortical neurons (**Fig. 4g, 4h, S4b, and S4c**). We observed no induction of type I IFN-induced ISGs upon treatment with IFN-beta in *IFNAR1* KO neurons compared to WT neurons, confirming impaired IFN signaling in *IFNAR1* KO cortical neurons at a functional level (**Fig. 4f and 4g**). When we inoculated WT and *IFNAR1* KO hPSC-derived cortical neurons with MPXV (MOI = 1), there was no increase in MPXV infected cells in *IFNAR1* KO neurons compared to WT neurons at early and late time points post infection (**Fig. 4h, S4e, and S4f**), suggesting that inducible IFNAR-dependent signaling is dispensable for neuronal anti-MPXV immunity. To test whether neuronal constitutive rather than inducible type I IFN signaling is essential for neuronal anti-MPXV immunity, we pre-treated WT cortical neurons with Ruxolitinib, a JAK-STAT inhibitor, after which we performed MPXV inoculation (MOI = 1). At 24 hpi, we observed no increase in MPXV-infected cortical neurons compared to DMSO-treated neurons (**Fig. 4h, S4g, and S4h**), indicating that neuronal constitutive anti-MPXV immunity is type I IFN-independent. To determine potential regulators of constitutive neuronal anti-MPXV signaling, we identified a subset of genes, such as *CCL25*, *SFTPD*, *NOS2*, and *CCL3L1*, from our targeted pathway analyses that were expressed in both WT and *IFNAR1* KO neurons, but not significantly expressed in either of the glial cell types in basal conditions (**Fig. 4i**). All but one of these candidate genes (IL12RB2) were independent of the JAK-STAT signaling pathway. Together, these data suggest that constitutive, JAK-STAT-independent, rather than inducible type I IFN signaling is essential for neuronal anti-MPXV immunity.

## Discussion

The rise of hPSC-based disease modeling allows for the study of emerging pathogens such as SARS-CoV-2 and MPXV in human, disease-relevant cells of the brain(Harschnitz and Studer, 2021; Bauer et al., 2022). Recent work using human organoid models of the gut(Watanabe et al., 2023), skin(Li et al., 2023b), and kidney(Li et al., 2023a) has established the tropism of MPXV in these respective organ systems, whereas the tropism of MPXV for human CNS cells has remained unclear(Chailangkarn et al., 2022; Mahabadi et al., 2024; Schultz-Pernice et al., 2023). Using previously established protocols(Ciceri et al., 2022; Guttikonda et al., 2021; Lendemeijer et al., 2024), we identified that hPSC-derived astrocytes and microglia are more susceptible and permissive to MPXV clade IIb infection than cortical neurons. This increased susceptibility and permissiveness of glial cells to MPXV infection at a cellular level coincides with an impaired type I IFN and innate antiviral response at a molecular level and can be rescued by tecovirimat and IFN-beta treatment.

The efficient replication of MPXV in both astrocytes and microglia is consistent with its ability to cause severe encephalitis. Cell type-specific susceptibility to viral infection and infection-induced pathology is an important feature of many neurotropic viruses leading to virus-specific clinical phenotypes(Harschnitz and Studer, 2021). Previous work has demonstrated a selective tropism of viruses for various CNS cell types, such as NPCs (ZIKV(Qian et al., 2016; Tang et al., 2016; Cugola et al., 2016; Garcez et al., 2016)), trigeminal neurons (HSV-1(Zimmer et al., 2018)), dopaminergic neurons (SARS-CoV-2(Chen et al., 2021)), the choroid plexus (SARS-CoV-2(Pellegrini et al., 2020; Jacob et al., 2020; Yang et al., 2021)), and microglia (Rubella virus(Popova et al., 2023)). Although we observed a comparable tropism of clade IIb MPXV for hPSC-derived astrocytes, microglia, and neurons *in vitro* and in *ex vivo* human brain tissue, we cannot exclude the possibility that other cell types such as oligodendrocytes or endothelial cells contribute to MPXV-induced neuropathology. Due to the limited availability of human brain tissue, specific regional differences within the brain in viral susceptibility or information on the route of viral entry could not be assessed. Analyses of autopsy brain samples from patients with mpox encephalitis, experimental *in vivo* studies, and follow-up studies using complementary models will further our understanding of the neuropathogenesis of mpox in the acute and post-acute phases of infection.

Our RNA-seq analyses demonstrated cell type-specific transcriptional responses to MPXV clade IIb infection that were associated with an increased susceptibility and permissiveness of glial cells to MPXV infection. Innate antiviral signaling pathways involved in the regulation of IFN production and response to IFNs were specifically down-regulated in astrocytes and microglia, but not in cortical neurons. Constitutive IFN signaling and the binding of IFNs to the IFN-ab receptor (IFNAR) that leads to the activation of intracellular signaling and downstream interferon stimulated gene (ISG) expression are essential for viral restriction. The downregulation of this response in glial cells is therefore likely associated with the efficient infection and replication of MPXV in glial cells in our *in vitro* models. Accordingly, treatment of hPSC-derived CNS cells with IFN-beta was sufficient to significantly reduce MPXV infection. Previous studies have demonstrated the efficiency of type I IFN treatment inhibiting VACV infection *in vivo*(Rodriguez et al., 1991). Therefore, IFNs may be of potential therapeutic use to treat mpox encephalitis, either as monotherapy or as combination therapy with other antivirals(Kalil et al., 2005; Wintergerst and Belohradsky, 1992).

Poxviruses have evolved multiple evasion strategies against the host innate immune response including the IFN system(Haga and Bowie, 2005; Fernández de Marco et al., 2010; Arndt et al., 2015). For example, poxviruses can inhibit inducible IFN signaling by expressing viral proteins that function as soluble forms of cytokine receptors(Symons et al., 1995; Xu et al., 2008). Our data in *IFNAR* KO cortical neurons, which are equally resistant to MPXV infection as their WT counterparts, would suggest that, at least for human neuronal cells, MPXV antiviral immunity is regulated by intracellular or constitutive antiviral signaling rather than inducible IFN signaling. Furthermore, pre-treatment of hPSC-derived cortical neurons with Ruxolitinib, a JAK-STAT inhibitor, similarly did not make neurons more susceptible to MPXV infection, implying that neuronal cell-intrinsic anti-MPXV immunity is independent of JAK-STAT signaling. These results support previous findings that unique host antiviral programs regulate the differential cell type-specific susceptibility within the brain to neurotropic viruses(Cho et al., 2013).

There are currently no approved treatments for severe MPXV infection or for mpox encephalitis. The antiviral drug tecovirimat (Tpoxx), developed and used for the treatment of VARV infection, is being used for mpox treatment based on historical data. Recent studies using the current strains of circulating MPXV have demonstrated effective antiviral effects both *in vitro* and *in vivo*(Warner et al., 2022; Frenois-Veyrat et al., 2022; Bojkova et al., 2022). Our data show that Tpoxx treatment of astrocytes resulted in a partial inhibition of viral infection, strongly reducing the number of infected cells in a concentration-dependent manner. Whether this would be sufficient to suppress viral replication in the brain *in vivo* is unclear, although animal studies have shown that Tpoxx can cross the blood-brain barrier. Recent studies in endemic regions of mpox demonstrated no reduction of mortality following Tpoxx treatment in patients suffering from severe mpox suggesting the urgent need for future anti-MPXV drug discovery. Furthermore, there have been several reports describing complicated mpox cases that are unresponsive to tecovirimat due to single amino acid changes in the MPXV protein F13, known to cause drug resistance(Smith et al., 2023; Karan et al., 2024). Alternative therapeutic strategies could be tested in our hPSC-based CNS model, which functions as an ideal platform for anti-viral discovery, as we previously demonstrated for SARS-CoV-2 infection(Yang et al., 2024).

A potential limitation of our study is the use of highly pure, mono-cultures of hPSC-derived CNS cells. While this allows for the careful dissection of cell-intrinsic responses to MPXV infection, it precludes the study of cell-nonautonomous interactions which can be important in antiviral signaling. The use of more complex co-culture systems, organoid models, and *in vivo* studies will allow for a better understanding of cell-cell interactions, paracrine signaling, and viral spread.

Overall, our study suggests a broad tropism of MPXV clade IIb throughout the human brain with a preferential susceptibility of glial cells, particularly astrocytes, as a key target for MPXV infection. Notably, in stark contrast to neurons, glial cells down-regulated essential innate antiviral signaling cascades upon viral infection. Furthermore, we showed that IFN and tecovirimat treatment could inhibit viral infection. Building on our hPSC-based models and data, future research could investigate the cell-nonautonomous signaling that occurs upon MPXV infection, and the potential risk of developing long-term neurological deficits.

## Acknowledgements

We thank the Genomics Facility, Light Imaging Facility, and Flow Cytometry Unit at Human Technopole and the BSL-3 facility of the diagnostic unit of the Viroscience department at Erasmus Medical Center for technical support. We are grateful to members of the Van Riel lab and Harschnitz lab for insightful discussions. We acknowledge funding from the Human Technopole (O.H. and J.D.V.). L.B. is supported by The Netherlands Organization for Scientific Research (XS contract number OCENW.XS22.2.045) and by a grant 2023 from the European Society of Clinical Microbiology and Infectious Diseases (Europäische Gesellschaft für klinische Mikrobiologie und Infektionskrankheiten) (ESCMID). N.P. is a PhD student within the European School of Molecular Medicine (SEMM). E.C. is supported by an EMBO postdoctoral fellowship (EMBO ALTF 418-2022). This work was supported by the Netherlands Organ-on-Chip Initiative, an NWO Gravitation project (024.003.001) funded by the Ministry of Education, Culture and Science of the government of the Netherlands, the ZonMW PSIDER program TAILORED (10250022110002) (S.A.K., F.M.S.D.V.), and by an Erasmus MC Human Disease Model Award (F.M.S.D.V., H.S.). D.V.R. is supported by fellowships from The Netherlands Organization for Scientific Research (VIDI contract 91718308). O.H. is supported by the Warren Alpert Foundation, the Brain & Behavior Research Foundation, and Fondazione Telethon.

## Author Contribution

Conceptualization: O.H. and D.v.R. Methodology: L.B. and S.G. Virus production: L.B., L.L., and B.E.V. Sequencing of viral stocks: L.S. and B.O.-M. MPXV infections: L.B. Cortical neuron differentiation experiments: S.G. and N.P. Microglia differentiation experiments: S.G. and N.P. Astrocyte differentiation experiments: L.B. K.L H.S. F.M.S.D.V. and S.A.K. designed and supervised astrocyte differentiation experiments. RT-qPCR experiments: S.G. RNA-seq and bioinformatics analyses: F.Z. and J.D.V. Primary human brain tissue: J.S. and C.D. Primary human brain tissue cultures: A.B. and Z.G. IFNAR1 KO generation: F.P. and S.G. Imaging and image analysis: L.B., S.G., N.P., and E.C. Writing—original draft: O.H. Writing—review and editing: L.B., S.G., D.v.R., J.D.V., and O.H. All authors discussed and analyzed the data. Supervision: D.v.R. and O.H.

## Material and Methods

### VeroE6 cell culture

VeroE6 (ATCC CRL 1586) cells were maintained in Dulbecco’s modified Eagle’s medium (DMEM; Lonza) supplemented with 10% fetal calf serum (FCS; Sigma-Aldrich), 10 mM HEPES (Gibco), 1.5 mg/ml sodium bicarbonate (Capricornus Scientific), 2 mM L-Glutamine (Capricorn Scientific), 100 IU/ml penicillin and 100 mg/ml streptomycin (Capricornus Scientific). Cells were grown at 37°C in 5% CO2 and cells were passaged at confluency with the use of phosphate-buffered saline (PBS) and trypsin-EDTA (0.05%) (Capricornus Scientific). Cells were routinely checked for mycoplasma.

### Human pluripotent stem cells

To assess the neurotropism of MPXV H9 (WAe009-A(Ciceri et al., 2022), WiCell) embryonic stem cells and WTC-11 (UCSFi001-A, Coriell) induced pluripotent stem cells were used. These lines were maintained in embryonic stem cell-qualified matrigel coated plates and grown in Essential 8 (Thermo Fisher) supplemented with E8 supplements and Penicillin/Streptomycin (Thermo Fisher).

### Viruses

MPXV was isolated from a swab from a pox lesion of a Dutch patient on VeroE6 cells. The isolate belonging to the clade IIb is available through the European Virus Archive Global Ref-SKU: 010V-04721. From initial virus stock, MPXV was passaged two times by inoculating a confluent layer of VeroE6 cells with a multiplicity of infection of 0.1-0.01 in Advanced DMEM/F-12 (Gibco) supplemented with 10 mM HEPES, Penicillin/Streptomycin and 2mM L-Glutamine and 100 µg/ml primocin (Invivogen). The cells were harvested after 3-4 days when full cytopathic effect was observed. Residual cells were scraped off the flask and the lysate was centrifuged 2000x*g* for 2 min. The supernatant was discarded and the pellet was resuspended in OptiMEM (Gibco) and subjected to three times freeze thaw cycles in a dry-ice ethanol bath. The lysate was diluted in 20 ml Opti-MEM and spun down. The cell debris was discarded and the supernatant was used for plaque assay and next-generation sequencing. All MPXV stock propagation and experiments were performed in a Class II Biosafety Cabinet under BSL-3 conditions.

### DNA extraction

For amplicon sequencing of MPXV stocks, 110 μl of sample material was centrifuged at 10,000 *xg* for 5 min and 100 μl of supernatant was treated with Turbo DNase (Invitrogen) for 30 min at 37 °C. Nucleic acids were extracted using the High Pure Viral Nucleic Acid Kit (Roche) according to the manufacturer’s recommendations.

### Amplicon sequencing and data analysis

Multiplex PCR was performed as described previously (https://www.protocols.io/view/monkeypox-virus-whole-genome-sequencing-using-comb-n2bvj6155lk5/v1). Sequencing libraries were prepared using the Ligation Sequencing Kit (SQK-LSK110) (Oxford Nanopore Technologies) with Native Barcode Expansion (EXP-NBD196) (ONT) and sequenced on a GRIDIon (ONT). Reads were basecalled with Guppy v6.0.1 in high accuracy mode, demultiplexed using Porechop v0.2.4 (https://github.com/rrwick/Porechop) and mapped against ON563414.3 using Minimap2 v2.17 (https://github.com/lh3/minimap2). NextClade v2.7.0 was used for sequence quality checks, clade assignment and mutation calling(Aksamentov et al., 2021). Consensus sequences were aligned using MAFFT v7.475(Katoh and Standley, 2013).

### RT-qPCR

To validate cortical neuron differentiations we collected RNA from differentiating cells at different stages, *e.g*. hPSC, DIV0, DIV5, DIV10, DIV15, DIV20 and DIV40. At each time point cells were lysed using 350 μl RLT buffer (Qiagen) and RNA was extracted according to manufacturer’s instructions (Qiagen). Equal amounts of RNA from each sample were used for cDNA synthesis with the kit Revertaid First Strand cDNA synthesis Kit (Thermo Scientific). The obtained cDNA was diluted to 2-4 ng/µl for subsequent qPCR with the Sybr Green (Applied Biosystems) method. qPCR primers have been selected from the Origene Website or designed with different bioinformatic tools (sequences are reported in **Table 1**). For the analysis, the 2^(-ΔCt) method was adopted and the expression of genes of interest was normalized to the PSMB2 housekeeping gene.

**Table 1:**
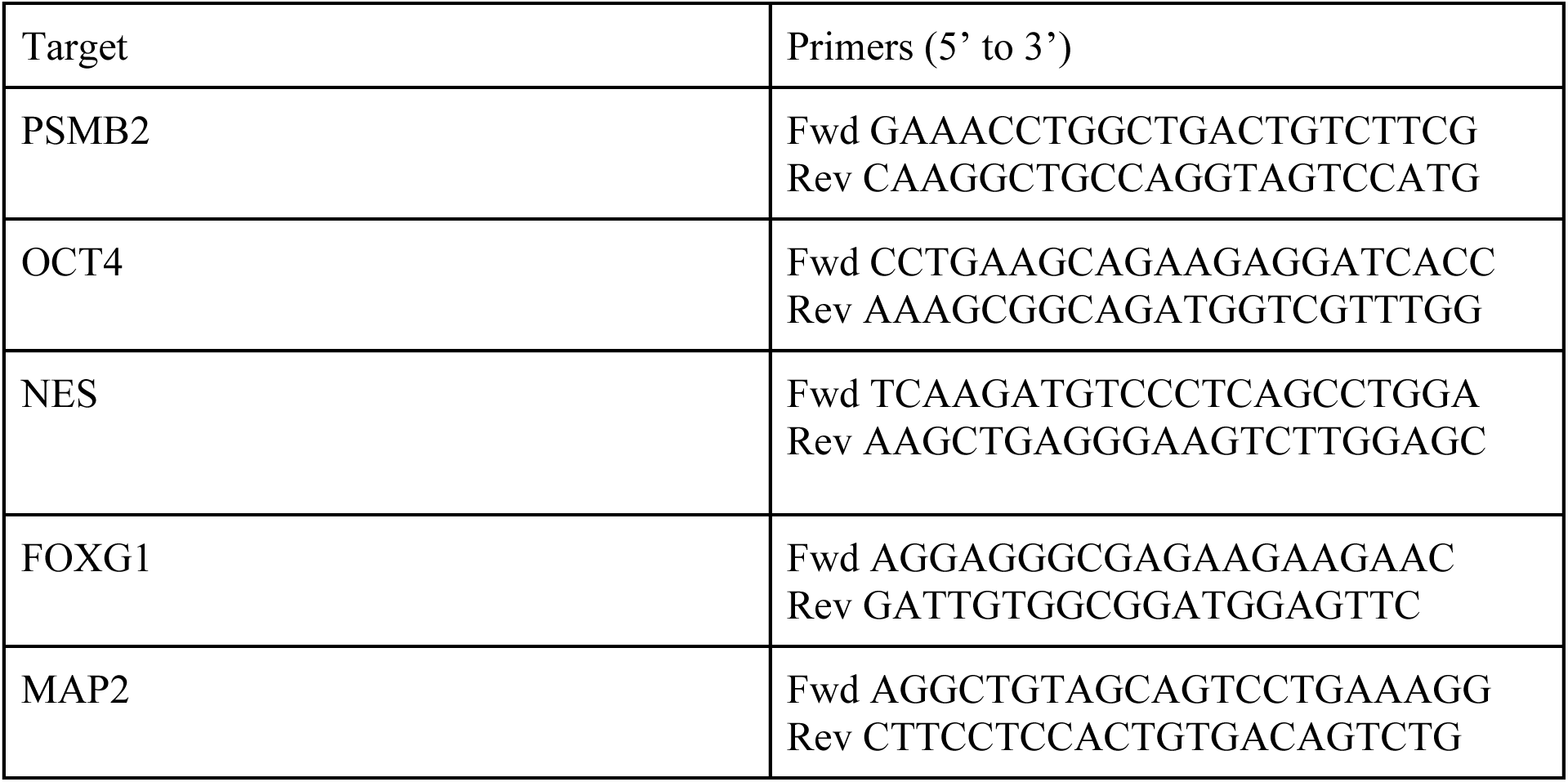
List of genes tested by RT-qPCR.

### Replication kinetics

Before infection of astrocytes, cortical neurons, and microglia, supernatant was removed and cells were infected with MPXV at an MOI = 1. As control, VeroE6 cells were infected with MPXV also at MOI = 1. After 1 h of incubation at 37°C, the inoculum was removed and astrocytes, microglia and VeroE6 cells were washed three times with PBS and fresh cell specific medium was added. In cortical neurons the inoculum was removed and without washing, fresh cortical neuron specific medium was added to prevent cell detachment. At the indicated time points, an aliquot of the supernatant was collected for subsequent analysis. All experiments were performed in biological triplicates.

### Plaque assay

For infectivity measurement, VeroE6 cells were seeded at 250.000 cells/well and incubated at 37°C in 5% CO2 overnight. Ten-fold serial dilutions of the collected supernatant in advanced DMEM/F-12 (Gibco) supplemented with 10 mM HEPES (Gibco), penicillin/streptomycin (Capricorn Scientific) and 2mM L-glutamine (Capricorn Scientific) were added to a monolayer of VeroE6 cells. After incubation at 37°C for 1 hour, the medium was taken off and cells were overlayed with 1.2% Avicel (FMC biopolymers) in Opti-MEM I (1X) + GlutaMAX. After 48 hours cells were fixed for 30 minutes in formalin and the whole plate was submerged in ice cold 100% ethanol. Cells were washed once with PBS followed by staining with 0.2% Crystal Violet (Sigma) in 20% Methanol (Sigma). Endpoint titers were calculated by visualizing plaques using light microscopy and plaque forming units/ml were calculated.

### Differentiation of cortical neurons

Cortical neuron differentiations were performed using a previously established protocol using WA-09 and WTC-11 hPSCs. Human PSCs were maintained on vitronectin (Thermo Fisher Scientific) at 37°C in 5% CO2 with Essential 8 medium (E8) and passaged once a week with EDTA. Prior to differentiation, hPSCs were dissociated into a single-cell suspension using Accutase and plated at 300,000 cells/cm2 onto geltrex-coated plates in E8 with ROCK inhibitor (R&D; Y-27632; 10 μM). Neural induction was achieved through dual-SMAD inhibition(Chambers et al., 2009) using Essential 6 Medium containing LDN193189 (Stemgent, 100 nM) and SB431542 (R&D, 10 μM), in combination with Wnt inhibition for 3 days. From day 11 onwards, medium was changed to a neural differentiation medium (1:1 DMEM/F12 and Neurobasal, 1× N2 supplement, 1× B27, 1× penicillin/streptomycin). At day 20, neural progenitor cells were either cryopreserved using Stem Cell Banker or replated onto poly-ornithine/laminin/fibronectin plates at 150,000 cells/cm2 in maturation medium. From day 21 onwards, media change occurred every 5 days until day 40, when cortical neurons were used for subsequent experiments.

### Differentiation of astrocytes

Human astrocyte differentiations were performed from NPCs that were made according to an embryoid body-based protocol(Gunhanlar et al., 2018), from the WA-09 and WTC-11 human PSC lines. In brief, NPCs were purified using fluorescence-activated cell sorting, according to a previously published protocol(Yuan et al., 2011). They were further differentiated to astrocytes through addition of 10 ng/ml BMP4 (Abcam) and 10 ng/ml LIF (Tebu Bio) to the NPC medium(Lendemeijer et al., 2024). The purity of astrocytes was assessed by GFAP and S100b staining.

### Differentiation of microglia

Microglia were differentiated according to a previously published protocol using WA-09 and WTC11 hPSCs(Guttikonda et al., 2021). In brief, primitive hematopoiesis was induced through a careful modulation of Wnt signaling. To achieve this, hPSCs were exposed on day 0 to the Wnt activator CHIR99021 (R&D, 3 µm), which was replaced with the Wnt inhibitor IWP2 (Tocris, 2 µm) after 18 hours. The quality of the differentiation was assessed before replating cells on day 3, when the expression of the CD235a marker of primitive hematopoiesis was quantified by flow cytometry (Table 1). Human PSC-derived hemangioblasts were provided the developmental cues needed to give rise to primitive erythromyeloid progenitors (EMPs) by day 10. The quality of the differentiation at this stage was tested by quantifying the expression of CX3CR1 by flow cytometry (Table 1). EMPs were subjected to M-CSF (R&D, 10 ng/ml) and IL34 (R&D, 100 ng/ml) to obtain mature microglia on day 30, when they were used for viral experiments. The quality of the differentiation was tested by immunofluorescence staining for the microglial marker IBA1.

### KO generation using CRISPR-Cas9

Single guide RNA sequences (sgRNA) were generated using the online design tool from Synthego (https://www.synthego.com/). We selected two sgRNAs targeting *IFNAR1* exon 2 (target specific region of selected gRNAs: 5′-AAAATCTCCTCAAAAAGTAG-3′ and 5′-TAGATGACAACTTTATCCTG-3′), and each was inserted into the px459-pSpCas9-2A-Puro after BbsI (Thermo Fisher) restriction enzyme treatment. The plasmid was then cloned in Top10 competent *E. coli* cells and purified for lipofection in H9 embryonic stem cells. Human PSCs were plated on Geltrex-coated plates in colonies of 2-3 cells one day before transfection in the presence of rock inhibitor and grown in mTeSR1 media supplemented with mTeSR1-growth factors and 100 U per ml penicillin, 0.1 mg per ml streptomycin (all Life Technologies) at 37 °C with 5% CO2. The plasmid bearing the guide RNA was then used to transfect H9 cells in the presence of Lipofectamine Stem Transfection reagent (invitrogen); complete medium change was performed the following day and after 48 hours 0.66μg/ml puromycin was added to the culture for 24 hours. Cells were then kept in mTeSR1 until colonies wells were 80-90% confluent. At that point cells were plated as single cells in matrigel-coated 96-well plates and grown in Stemflex media with supplements, CloneR (added upon plating and at the first medium change, Stem Cell Technologies), and penicillin, 0.1 mg per ml streptomycin (all Life Technologies) at 37 °C with 5% CO2 for about 10 days. Colonies were then picked for expansion and Sanger sequencing. For *IFNAR1*-KO genotyping, the following primers were used in this study (forward for guide 1, 5′-CTGTGCTGGGAGCAATCATTAG-3′; forward for guide 2, 5′-TCAGGGATGTGAGGGATAGAATAAC-3′; reverse, 5′-GATGTATTTGACTAAGTCTCTACAG-3′). To assess the pluripotency of the *IFNAR1* KO clones generated from this process, cells were stained for the markers NANOG and OCT4 according to the protocol described.

### Immunofluorescence staining

Cells were fixed with 10% formalin for 30 minutes at room temperature. Cells were permeabilized with 1% Triton-X (Sigma Aldrich) in PBS for 15 minutes on room temperature followed by a blocking step with blocking buffer (PBS supplemented with 1% BSA (Sigma Aldrich) and 0.5% Triton-X (Sigma Aldrich)) for 30 minutes on room temperature. The SOX2 antibody was pre-conjugated overnight with Zenon™ Rabbit IgG Labeling Kits Alexa-647 (Invitrogen, Z25308). Primary antibody incubation was performed overnight at room temperature in blocking buffer. Cells were washed three times in PBS and secondary antibody incubation was performed in blocking buffer for 1 hr at room temperature. To visualize cell nuclei, cells were incubated with a solution of Hoechst (Invitrogen, H3570) which was added during secondary antibody incubation. Slides were washed in PBS, dipped in water and mounted in ProLong Antifade Mountant (Thermo Fisher). Samples were imaged using a Zeiss LSM 700 confocal microscope.

**Table 1.** Antibodies used for stainings.

| Antibody | Dilution | Manufacturer | Cat # | Lot |
| --- | --- | --- | --- | --- |
| MAP2 | 1:200 | Synaptic Systems | 188004 | 30021200 |
| MAP2 | 1:1000 | Thermo Fisher | PA1-10005 |  |
| CTIP2 | 1:500 | Abcam | ab18465 |  |
| S100b | 1:200 | Sigma-Aldrich, | S2532 |  |
| GFAP | 1:200 | BD Pharma | 556327 | 2277814 |
| GFAP | 1:100 | Invitrogen | 14-9892-82 |  |
| Vaccinia Virus<br>Lister-FITC | 1:1000 | Abbexa | abx022301 |  |
| IBA-1 | 1:200 | Abcam | AB5076 |  |
| IBA-1 | 1:1000 | Wako | 019-19741 |  |
| HuC/HuD | 1:1000 | Abcam | ab184267 |  |
| NANOG |  | Abcam | ab109250 |  |
| OCT4 |  | Abcam | ab19857 |  |
| CD235a - APC | 1:400 | BD | 551336 |  |
| CX3CR1 - 421 | 1:100 | Biolegend | 341619 |  |
| donkey anti<br>mouse AF555 | 1:1000 | Invitrogen | A31570 | 2153958 |
| donkey anti<br>mouse AF647 | 1:1000 |  | A31571 | 2311820 |
| goat anti guinea<br>pig AF647 | 1:1000 |  | A21450 | 2110845 |

### Patient material for human brain slices

Damaged human brain tissue was used after approval of the Medical Ethical Committee of the Erasmus University Medical Center (protocol MEC-2022-0776). Written informed consent was obtained prior to surgery in accordance with the Declaration of Helsinki. Tissue samples were obtained from patients undergoing brain tumor surgery at the Department of Neurosurgery (Erasmus University Medical Centre Rotterdam, the Netherlands). Infiltrated peri-tumoral neocortical specimens were removed during the regular course of tumor resection. Human patient samples were anonymized manually.

### Acute brain slice preparation

Immediately after resection, human peritumoral cortex was placed in ice-cold oxygenated (95%O2/5%CO2) artificial cerebrospinal fluid (aCSF) containing (in mM): 124 NaCl, 24 NaHCO3, 22 D-glucose, 2.5 KCl, 1.25 NaH2PO4, 2 MgSO4, and 2 CaCl2. 350 um thick coronal slices were prepared on a vibratome (VT1200S, Leica) in oxygenated (95%O2/5% CO2) ice-cold N-methyl-d-glucamine (NMDG)-based slicing solution containing (in mM): 93 NMDG, 93 HCl, 25 D-glucose, 30 NaHCO3, 2.5 KCl, 0.5 CaCl2, 10 MgCl2, 20 HEPES, 1.25 NaH2PO4, 12 N-acetyl-cysteine, 2 thiourea, 5 Na-ascorbate, 3 Na-pyruvate (300 mOsm, pH 7.4).

### Infection of human brain slices

350 micron thick brain slices from the cortical regions were transferred into 12-well trans-well inserts (Corning). The apical compartment and the basolateral compartment were filled with cortical neuron medium. The brain slices were infected with an equivalent of 6*10^6 PFU of MPXV in the apical compartments. After 1 hour of incubation, brain slices were washed once with PBS and fresh medium was added. 24 hrs after infection, supernatant was removed and wells were filled with 10% formalin and cells were fixed for 3 days at room temperature.

### Multiplex Fluorescent RNAscope® and immunofluorescence staining of *ex vivo* human brain slices

After fixation, brain slices were transferred in PBS with 30% sucrose until fully immersed, embedded in OCT compound (Scigen), frozen rapidly, and stored at −20°C. 15 micrometre-thick sections were sliced on a cryostat (Leica). To detect MPXV mRNA, RNAscope® using Multiplex Fluorescent V2 Assay (ACD, Cat. No. 323100) was integrated to immunofluorescence staining in a workflow optimized for RNA and protein detection. Both mock and infected brain slices were probed using the RNAscope® Probe-V-Monkeypox (ACD, Cat. No. 534671). The protocol followed ACD manufacturer’s instructions, and all the reagents used for the RNAscope® belong to the RNA-Protein Co-Detection Ancillary Kit (ACD, Cat. No. 323180). Briefly, after washing the slides with PBS1x, tissue was dehydrated by EtOH gradient (50%, 70%, 100%), and treated with RNAscope® Hydrogen Peroxide for 15 minutes at RT. Target retrieval was performed using a steamer with the RNAscope® 1x Co-Detection Target Retrieval Reagent for 15 minutes, and primary antibodies were diluted in Co-Detection Antibody Diluent and incubated at 4°C overnight. Primary antibodies used are Mouse α GFAP (Invitrogen), Rabbit α IBA1 (Wako), Rabbit α HuC/HuD (Abcam), see Table 1. Following primary antibody incubation, sections were post-fixated with 10% Neutral Buffered Formalin (Bio-Optica, 05-01004F) for 30 minutes at RT, treated with RNAscope® Protease Plus at 40°C for 30 minutes in the HybEZ^TM^ Humidifying System, and probed with the RNAscope® Probe-V-Monkeypox for 2 hours at 40°C in the HybEZ^TM^ oven. Probe signal was amplified by consecutive hybridization of AMP1 (30 minutes at 40°C), AMP2 (30 minutes at 40°C), and AMP3 (15 minutes at 40°C), developed using the RNAscope® Multiplex FL v2 HRP-C1 (15 minutes at 40°C) and the TSA Vivid Fluorophore 650 (used 1:1500 diluted in TSA buffer, for 30 minutes at 40°C, ACD, Cat. No. 323273). After a blocking step with the RNAscope® Multiplex FL v2 HRP blocker (15 minutes at 40°C), secondary antibodies were diluted in Co-Detection Antibody Diluent and incubated at 40 minutes at RT. Secondary antibodies used are 1:1000 Donkey α Mouse Alexa Fluor™ Plus 555 (Invitrogen, A32773) or 1:1000 Donkey α Rabbit Alexa Fluor™ Plus 555 (Invitrogen, A32794). After the DAPI counterstain, the slides were mounted with ProLong™ Glass Antifade Mountant (Thermo Fisher, P36984). To visualise MPOX infection in the tissue, the slides were imaged by confocal microscope LSM 980 (Zeiss), with objective magnification of 63x, (16 bits, frame size 2048 pixels x 2048 pixels).

Immunofluorescent staining was performed to characterize the cellular composition of the brain tissue (**Fig. S2m and S2n**). Brain slices were rinsed in PBS1x, incubated with Retrieval Solution (Dako) for 45 minutes at 70°C, permeabilized with 0,5% Triton X-100 in PBS1x for 20 minutes, and blocked with 0,1% Triton X-100, 10% Donkey Serum in PBS1x for 1 hour at RT. Afterwards the brain sections were incubated with primary antibodies diluted in blocking solution at 4°C overnight. After washing, primary antibodies were visualized with 1:1000 Donkey α Mouse Alexa Fluor™ Plus 555 (Invitrogen, A32773) or 1:1000 Donkey α Rabbit Alexa Fluor™ Plus 647 (Invitrogen, A31573). Nuclei were stained with DAPI (Thermo Fisher, 62248). Immunostained sections were scanned using confocal microscope LSM 980 (Zeiss), with objective magnification of 40x (16 bits, frame size 2048 pixels x 2048 pixels).

### Tecovirimat rescue experiments

Multi-cycle MPXV infection was performed by incubating a confluent layer of hPSC-derived astrocytes at a MOI 1 for 1 hour at 37 °C. Next, supernatant was removed and cells were washed three times with PBS. After the washing steps, fresh astrocyte medium with a serial dilution of tecovirimat (Selleckchem) was added to the cells. At the indicated timepoints, medium was collected to determine virus titers by plaque assay, cells were fixed to determine the infection percentage by immunofluorescence staining, or cells were lysed for RNA isolation at indicated timepoints. Tecovirimat was dissolved in DMSO at 10mM stock concentration and stored at - 20°C. Cell viability was measured with an CellTiter-Glo® Luminescent Cell Viability Assay (Promega) and values were compared to mock treatment. The concentration of compound that inhibits plaque forming units by 50% (IC50) was calculated by nonlinear regression analysis with GraphPad Prism Version 10.

### Interferon-beta and Ruxolitinib treatment

To assess the protective role of IFN-beta on CNS cells, cortical neurons and astrocytes were treated with 10 ng/ml (100U) IFN-beta (Peprotech) for 24 hours before infection with MPXV (MOI 1). To assess the relevance of IFN signaling in protecting against MPXV infection, astrocytes and cortical neurons were treated for 24 hours before infection with 8 µM of Ruxolitinib (reconstituted in DMSO to a stock concentration of 10 mM, SelleckChem), a JAK-STAT inhibitor. Cells were fixed 24 hours after MPXV infection and stained according to the already described protocol.

To assess baseline and type I interferon-induced signaling, cortical neurons from wild type and *IFNAR1* KO (clone #1) background were differentiated following the protocol described above. DIV40 cortical neurons were treated with or without 10 ng/ml (100U) IFN-beta (Peprotech) for 4 hours and samples were lysed in 350 μl RLT buffer prior to sequencing.

### RNA sequencing

Infections were performed at an MOI of 1 and samples were harvested 24 hpi. Total RNA was extracted using the RNeasy Mini kit (Qiagen) following the on column DNAse digestion protocol, according to manufacturer’s instructions. RNA-seq libraries of polyadenylated RNA were prepared using Illumina Stranded mRNA Prep (Illumina) according to the manufacturer’s instructions and sequenced on an NovaSeq 6000 PE100 (Illumina).

Paired-end reads were aligned to the human genome reference GRCh38.p13 with the R software package Rsubread(Liao et al., 2019). Alignment BAM files were merged using Samtools (1.14-python-3.9.10) when needed. Mapped reads were summarised to gene level counts with the featureCounts function of Rsubread, using reference gene annotations (v.42) downloaded from the GENECODE project (https://www.gencodegenes.org/human/). Protein coding genes with detected counts in at least one sample library were retained and normalized using TMM normalization. For viral read mapping the same procedure was followed, using the Monkeypox virus genome (GenBank No.ON563414.3) and annotation as reference. Instead of TMM normalization, viral reads were converted to TPM.

#### Differential expression and pathway enrichment analysis

Differential expression analysis was performed with the edgeR package. Genes with a FDR-corrected P < 0.01 and an absolute log2 fold change (FC) > 1.0 were considered as differentially expressed. Differential expression analysis results for all cell types are reported in Supplementary Tables. Gene ontology and functional enrichment analyses were performed using Fast Gene Set Enrichment Analysis (fgsea) for ranked gene lists and Enrichr (https://maayanlab.cloud/Enrichr/) and gprofiler2 (https://cran.r-project.org/web/packages/gprofiler2/index.html) for gene sets. Statistical analyses and plots were performed using the programming language R (R Core Team, 2012). Gene ontology biological processes (GO:BP), KEGG, and REACTOME databases were used as knowledge base for gene set enrichment analyses.

#### Pathway activity scores and differential pathway activity analysis

Pathway activity scores were calculated for each RNAseq sample by gene set variation analysis as implemented in the R package oppar (v.1.32.0), using the gsva() function. Briefly, GSVA estimates a normalized relative expression level per gene across samples. This expression level is then rank-ordered for each sample and aggregated into gene set or pathway scores by calculating sample-wise enrichment using a Kolmogorov–Smirnov-like rank statistic. Antiviral related pathways (GO:0035455, GO:0035456, GO:0034341, GO:0034340, GO:0032647, GO:0032648, GO:0032649, GO:0032479, GO:0034097, GO:0001817, GO:1990868, GO:0032642) were extracted from GO Biological Processes v2023 downloaded from https://maayanlab.cloud/Enrichr/#libraries, considering terms involving innate immunity, interferon, interleukin, cytokine, and chemokine. Viral related pathways (GO:0044827, GO:0044788, GO:0044828, GO:0044793, GO:0043922, GO:0045869, GO:0046597, GO:0045071, GO:1903901, GO:0048525, GO:0032897, GO:0039532, GO:0044829, GO:0044794, GO:0043923, GO:0046598, GO:0045070, GO:1903902, GO:0048524, GO:0045091, GO:0046596, GO:0045069, GO:1903900, GO:0046782, GO:0039531, GO:0044790, GO:0046784) were extracted by filtering Gene Ontology biological processes containing the keyword “viral.” Differential pathway activity analysis was performed for each pathway-cell type combination by testing for an association between pathway activity scores and infection status using a linear model and computing t-statistics as implemented in the lmfit() and eBayes() functions from the Limma R package (v.3.60.6).

### Statistical analysis

Statistical analysis was performed as explained in detail in the figure legends with the latest available version of GraphPad Prism.

---

**Supplementary Fig. 1:**
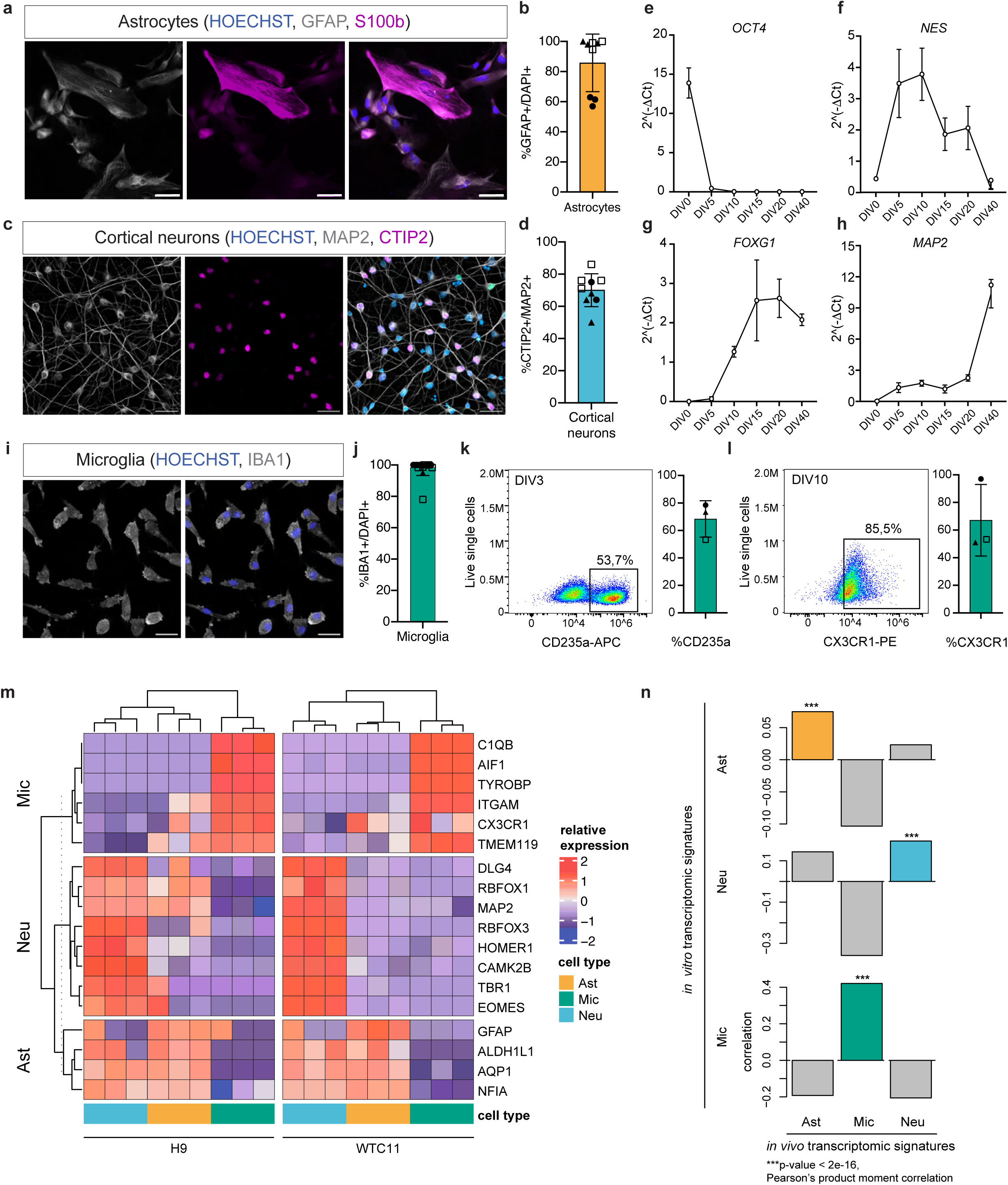
Characterization of hPSC-derived astrocytes, cortical neurons, and microglia. **a**, Mature astrocytes at DIV65 were stained for the canonical astrocytic markers GFAP (grey) and S100b (magenta). Scale bar = 20µm. **b**, Quantification of GFAP+/DAPI+ cells at DIV65 over 3 biological replicates. **c,** Mature cortical neurons at DIV40 were stained for MAP2 (grey) and the cortical layer V marker CTIP2 (magenta). Scale bar = 50 um **d**, Quantification of CTIP2+/DAPI+ cortical neurons at DIV40 over 3 biological replicates. **e-h**, Characterization of the cortical differentiation by reverse transcription quantitative polymerase chain reaction with stem cell markers (**e, f**) and neuronal markers (**g, h**). **i**, Characterization by flow cytometry of three independent microglia differentiation at DIV3 with the erythrocyte progenitor marker CD235a. **j**, Characterization by flow cytometry of three independent microglia differentiation at DIV10 with the microglia marker CX3CR1. **k,** Mature microglia at DIV30 were stained for the canonical microglial marker IBA1 (grey). **l,** Quantification of IBA1+/DAPI+ cells at DIV30 over three biological replicates. Scale bar = 20 um. **m**, Relative expression profiles for cell type marker genes in mock-infected WA-09-derived CNS cells (n = 3 independent samples per condition). **n,** Distribution of transcriptomic similarity values (signature correlation) comparing in vivo transcriptomic signatures(Zhang et al., 2016) to our WA-09-derived CNS cell transcriptional signatures. **a-b**, Experiments were performed using WTC11-derived astrocytes. **c-j**, Experiments were performed using WA-09-derived cortical neurons and astrocytes. **b**, **d, l,** *N* = 3 independent biological replicates, indicated by different shapes. *n* = 9-12 technical replicates. **i, j,** *N* = 3 independent biological replicates, indicated by different shapes. *n* = 2 technical replicates averaged for each data point. Data represent mean ± SD from three pooled experiments. All experiments were performed in two to three biological replicates

**Supplementary Fig. 2:**
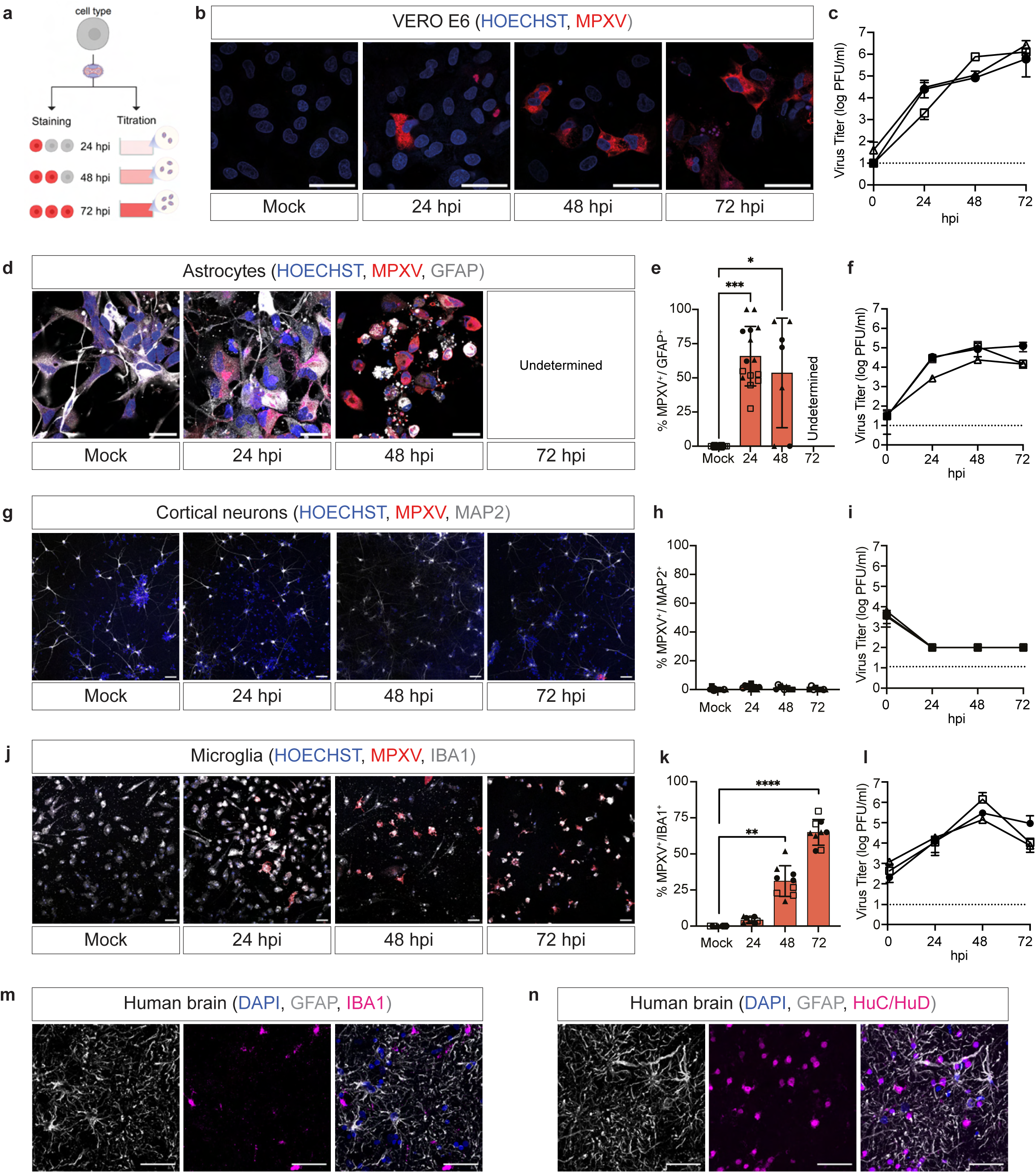
Astrocytes, cortical neurons, and microglia derived from WTC-11 background are differentially susceptible and permissive to MPXV infection. **a**, Schematics representing experimental design. Astrocytes, cortical neurons and microglia (represented for simplicity by a round grey cell) were infected with MPXV at 1 MOI. The susceptibility of each cell was assessed by immunostaining over three timepoints (24, 48 and 72 hours post infection [hpi]). The permissiveness was measured by titrating viral particles released in the supernatant over the aforementioned timepoints. **b-c**, Immunofluorescence staining and titration assays of VeroE6 cells (positive control) infected with a multiplicity of infection of 1 as positive control tested by immunofluorescence staining (**b**) and titration assays (**C**), both showing an increased amount of MPXV over time. **d-e,** Immunofluorescence staining for astrocyte marker GFAP (grey) and MPXV antigen (red), with quantification (**e**) of MPXV+/GFAP+ cells. Images representative of three independent experiments. **f,** Plaque forming assays of supernatant collected from MPXV-infected astrocytes. **g-h,** Immunofluorescence staining for neuronal marker MAP2 (grey) and MPXV antigen (red) with quantification (**h**) of MPXV+/MAP2+ cells. Images representative of three independent experiments. **i,** Plaque forming assays of supernatant collected from MPXV-infected cortical neurons. **j-k,** Immunofluorescence staining for microglial protein IBA1 (grey) and MPXV antigen (red) with quantification (**k**) of MPXV+/IBA1+ cells. Images representative of three independent experiments. **l,** Plaque forming assays of supernatant collected from MPXV-infected microglia. **m-n**, Immunofluorescence staining of human damaged brain tissue for astrocyte marker GFAP (gray), cortical neruon marker HuC/HuD (magenta, right panel) and microglia marker IBA1 (magenta, left panel). **d-l**, Experiments were performed using WTC11-derived astrocytes, cortical neurons and microglia. d-l, *N* = 3 independent biological replicates, indicated by different shapes. *n* = 7-15 technical replicates. Statistical analysis of image quantifications was performed using one-way ANOVA with Kruskal Wallis test. *p < 0.05, **p < 0.01, ***p < 0.001, ****p < 0.0001. All data represent mean ± SD. All experiments were performed in two to three biological replicates

**Supplementary Fig. 3:**
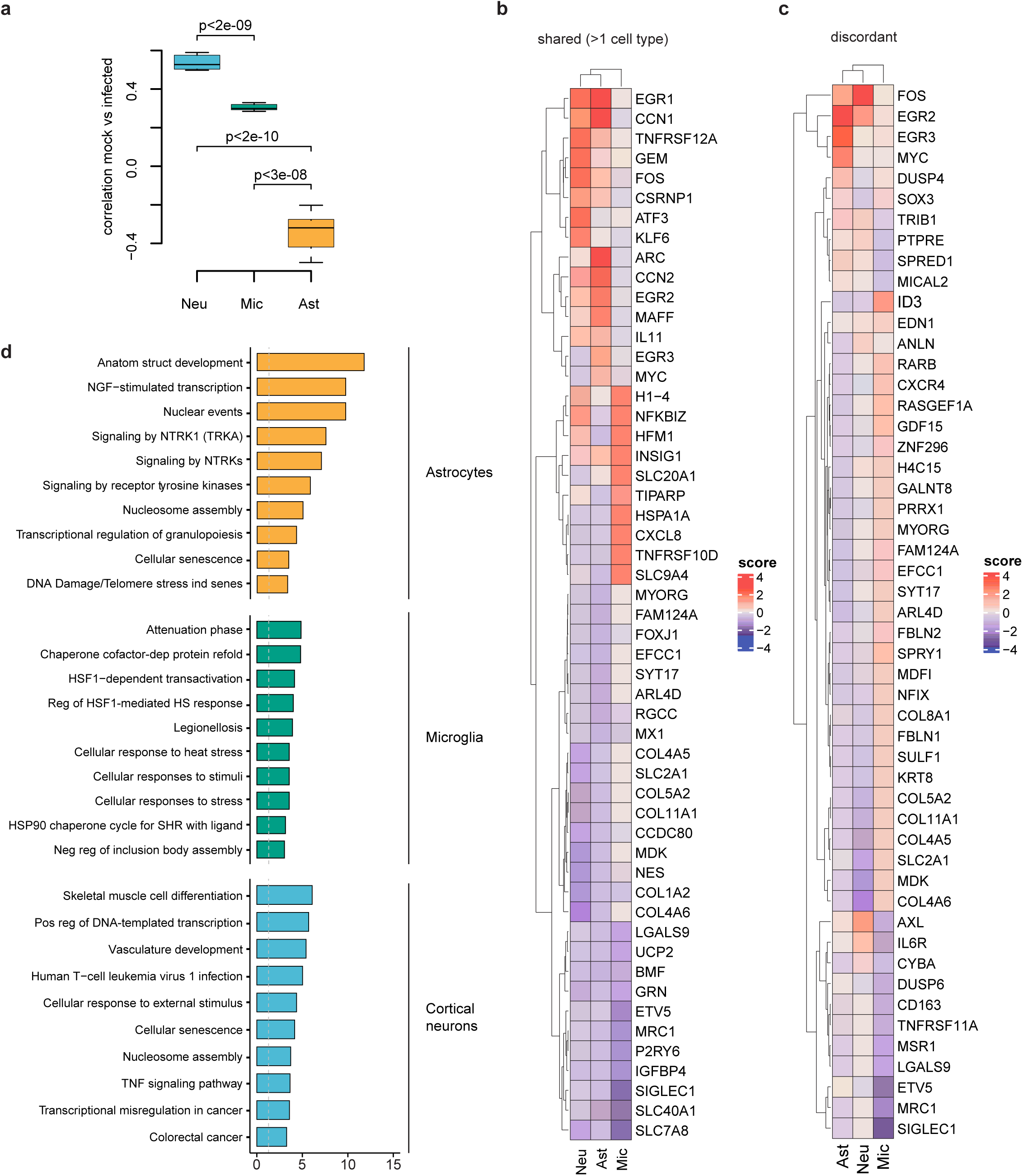
Transcriptomic changes in MPXV infected CNS cells. **a**, Distribution of transcriptomic similarity values (signature correlation) between mock- and MPXV-infected samples by cell type. **b-c**, Relative expression profiles for genes with a shared (**b**) pattern amongst at least two cell types, or a discordant (**c**) expression pattern. **d**, Upregulated pathways in astrocytes, microglia and cortical neurons after MPXV infection. **a-d**, experiments were performed using WA-09-derived astrocytes, cortical neurons and microglia.

**Supplementary Fig. 4:**
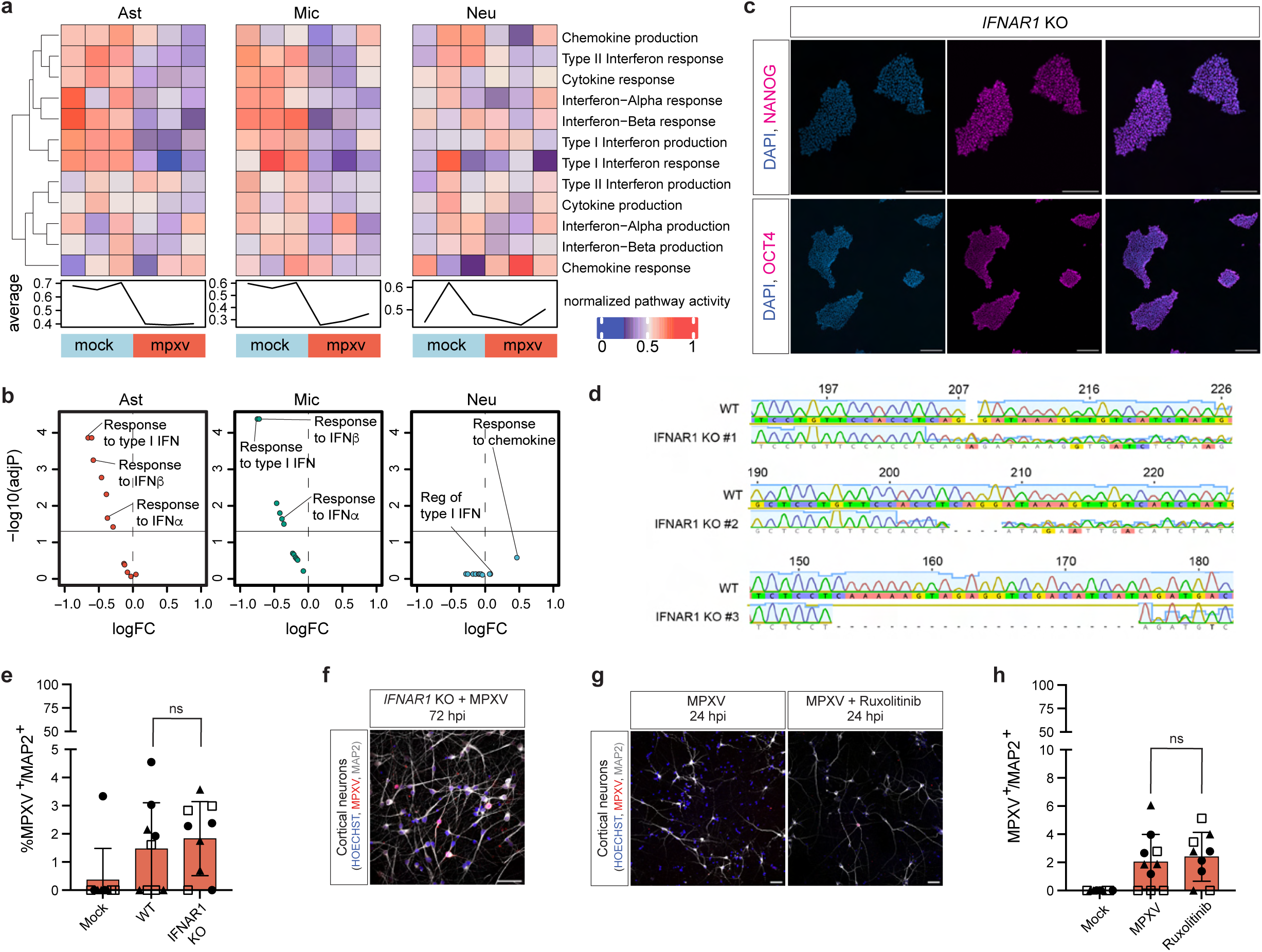
*IFNAR1* KO cortical neurons show similar infection to wild type cortical neurons. **a**, Pathway activity scores for interferon-associated GO processes in the WTC-11 background. Scores were computed per sample within each cell type. Bottom line plot depicts average per sample activity of all antiviral processes (rows). **b**, Volcano plot depicting differential pathway activity analysis of interferon-associated GO processes in mock- vs MPXV-infected WTC-11 hPSC-derived astrocytes, microglia, and cortical neurons (adjusted p < 0.05, linear model, mock- versus MPXV-infected). **c**, Representative immunofluorescence staining of *IFNAR1* KO clone #3 showing the expression of stem cell-related markers (NANOG, OCT4, depicted in magenta) after CRISPR-Cas9 editing. Scale bar= 50 um. **d**, Scheme of the mutations of the clones used in this manuscript induced by CRISPR-Cas9 editing. **e**, Quantification of the percentage of MPXV+/MAP2+ cortical neurons in the wild type and IFNAR1 KO (clones #1, #2, #3) background at 24 hpi. **f**, Representative image of *IFNAR1* KO cortical neurons at 72 hpi. **g-h**, Immunofluorescence staining and quantification of MPXV+/MAP2+ cortical neurons derived from WTC11 hPSCs. Images show, from left to right, untreated MPXV-infected cortical neurons and Ruxolitinib (JAK-STAT inhibitor) pre-treated MPXV-infected cortical neurons. Images representative of three independent experiments. **e-f,** Experiments were performed using WA-09-derived cortical neurons. **g-h,** Experiments were performed using WTC11-derived cortical neurons. **e, h,** *N*=3 independent biological replicates, indicated by different shapes. *n* = 9 technical replicates. Statistical analysis of image quantifications was performed using one-way ANOVA with Kruskal Wallis test. *p < 0.05, **p < 0.01, ***p < 0.001, ****p < 0.0001. All data represent mean ± SD. All experiments were performed in two to three biological replicates

